# Why linkage disequilibrium measures disagree: Fisher geometry of rare–common haplotype structure

**DOI:** 10.64898/2026.07.02.736022

**Authors:** Yuki Ichikawa

## Abstract

Conventional LD measures such as r^2^ perform poorly in the rare–common regime, particularly in asymmetric configurations such as nested haplotype structure. Because r^2^ is symmetric and quadratic, it removes directional structure in two ways: squaring discards the sign, or phase, retained by the signed LD coefficient D, while symmetric normalization hides the asymmetry between the conditional probabilities P(A|B) and P(B|A). Although D recovers the phase, it is locus-symmetric and unnormalized; its magnitude is hard to compare across frequency regimes and it does not by itself express which way the asymmetry runs.

We therefore analyze the conditional-probability asymmetry Δ = P(A|B) − P(B|A), together with r^2^ and D, as distinct scalar functions on the haplotype simplex under the Fisher information metric. The conditional probabilities P(A|B) and P(B|A) are bounded in [0, 1], directly express carrier-set inclusion, and are more readily visualized than D. Moreover, their difference admits the exact decomposition Δ = M + C into a marginal-frequency term M and an LD-coupled term C.

Prior work has characterized either the mathematical behavior of LD normalizations across allele-frequency space or the Fisher geometry of the haplotype simplex, but not their connection. We bridge this gap by showing that the geometric structure of the simplex explains why LD measures disagree in the rare–common regime and why symmetric normalizations such as r^2^ lose directional information. We show that the fixed-frequency leaf is intrinsically anisotropic, positively curved, and frequency-dependent under the Fisher metric. These geometric predictions are tested empirically — in phased 1000 Genomes data^1^ and a two-locus Wright–Fisher model — in a companion paper (Ichikawa, preprint^2^); the present note develops the geometry itself.

## 1. Introduction

### Why conventional LD measures disagree

The quantification of linkage disequilibrium (LD) between biallelic loci has relied, for more than half a century, on a family of related scalar statistics derived from the two-locus haplotype frequency distribution. The LD coefficient D = x_AB − p_A p_B (Lewontin and Kojima)^3^, the normalized D′ = D / D_max (Lewontin 1964)^4^, the squared correlation r^2^ = D^2^ / (p_A q_A p_B q_B), and various alternative normalizations (Hedrick 1987)^5^ remain in routine use. Each is defined as a different function of the same underlying haplotype frequencies, and each is reported as a single scalar value. In applications such as GWAS tagging, fine-mapping, and cross-population LD comparison, these scalars are routinely compared and thresholded as if their numerical distances were directly comparable across frequency regimes.

The disagreement among measures is most pronounced in the rare–common frequency regime. For example, two loci in apparent complete positive LD with *p*_*A*_=0.01 and *p*_*B*_=0.30 yield *D*=0.007, *D*^’^=1.000, *r*^2^=0.024, and an LD-coupled asymmetry component ∣*C*∣=0.68: four apparently inconsistent summaries of the same haplotype distribution. Each value is correct given its definition. The question is therefore not which measure is right, but why correct measures disagree, and what that disagreement implies for which structures remain visible under each normalization.

This limitation is especially evident in asymmetric haplotype configurations, such as nested structure, where one variant is preferentially carried on the background of another. Because *r*^2^ is symmetric and quadratic, it suppresses the asymmetry between *P*(*A*∣*B*) and *P*(*B*∣*A*). In the rare– common regime, this suppression becomes structural: when the rarer allele is nested within the commoner background, the bound *r*^2^≤*p*_*A*_*q*_*B*_/(*q*_*A*_*p*_*B*_) ensures that no configuration with *p*_*A*_≪*p*_*B*_ can achieve high *r*^2^, even under complete positive LD. The rare-variant signal is therefore not merely hard to detect under *r*^2^; it is geometrically compressed by the normalization itself.

### Directional asymmetry: the conditional-probability difference *Δ*=*P*(*A*∣*B*)-*P*(*B*∣*A*)

The problem is not only that *r*^2^ loses sign. Because it is also locus-symmetric, it cannot distinguish *P*(*A*∣*B*) from *P*(*B*∣*A*). The signed coefficient *D* restores coupling versus repulsion, but carries phase rather than direction: it remains symmetric in the two loci and unnormalized across frequency regimes. We therefore work directly with *P*(*A*∣*B*) and *P*(*B*∣*A*), which are bounded in [0,1]and directly express carrier-set inclusion. Their difference,

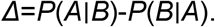

makes the missing direction explicit and separates exactly into marginal-frequency and LD-coupled components, as introduced in Section 2

### Fisher geometry of the haplotype simplex

Several lines of prior work have addressed aspects of this problem. Kang and Rosenberg (2020) systematically characterized LD statistics defined as normalizations of *D*, including their bounds, mean maxima, and accessible regions across allele-frequency space. Tanaka ^6^ proposed an equal-tempered LD measure designed to reduce frequency dependence. On a more geometric level, Shahshahani (1979)^7^ showed that the natural gradient on the allele-frequency simplex coincides with the Fisher-metric gradient, and Hofrichter, Jost, and Tran ^8^ established that the haplotype simplex under the Fisher information metric is a positively curved manifold, specifically the positive octant of a 3-sphere, with linkage equilibrium corresponding to a product-of-spheres structure.

The present work connects these lines of thought. We treat the Fisher-curved haplotype simplex as the natural geometric setting in which LD statistics are scalar functions of the same underlying haplotype distribution. This framing turns a familiar empirical difficulty — the dramatic disagreement among *r*^2^, *D, D*^’^, and ∣*C*∣for rare–common variant pairs — into a geometric question: which directions and regions of the simplex are expanded, compressed, or collapsed by each normalization? In this view, disagreement among LD measures is not merely statistical noise or imperfect estimation, but a consequence of applying different scalar projections to a curved, frequency-dependent space.

### The present study

Here we analyze the two-locus haplotype simplex under the Fisher information metric, treating *Δ*=*P*(*A*∣*B*)-*P*(*B*∣*A*), *r*^2^, and *D* as distinct scalar functions on the same curved space. We first characterize the fixed-frequency (*M,D*) leaf and show that its Fisher geometry is anisotropic, positively curved, and frequency-asymmetric. This provides a geometric explanation for why standard LD measures diverge most strongly in the rare–common regime, and why symmetric normalizations such as *r*^2^ obscure directional carrier-set structure.

These geometric predictions are evaluated empirically — in phased 1000 Genomes Phase 3 data and in a two-locus Wright–Fisher model — in a companion paper ^2^, which tests whether fixed r^2^ bins preserve information about nested versus non-nested haplotype configurations and whether the rare–common blind spot of r^2^ is structural rather than statistical. The present note develops the geometry on which those tests rest.

The paper is organized as follows. Section 2 introduces the conditional-probability asymmetry *Δ* and its decomposition into *M*+*C*. Sections 3–5 develop the Fisher geometry of the (*M,D*) and (*M,C*) representations. Section 6 interprets standard LD measures as scalar functions on this manifold. Section 7 discusses implications for rare-variant analysis and cross-population LD comparison. The empirical tests of these geometric predictions — in 1000 Genomes data and a two-locus Wright–Fisher model — are reported in a companion paper^2^.

## 2. The conditional-probability asymmetry Δ = P(A|B) − P(B|A) and its decomposition

For two biallelic loci with allele frequencies *p*_*A*_ and *p*_*B*_, haplotype frequencies *x*_11_=*f*_*AB*_, *x*_10_=*f*_*Ab*_, *x*_01_=*f*_*aB*_, *x*_00_=*f*_*ab*_, and LD coefficient *D*=*x*_11_-*p*_*A*_*p*_*B*_, we define the conditional-probability asymmetry *Δ*=*P*(*A*∣*B*)-*P*(*B*∣*A*).

Since

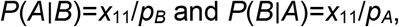

we have

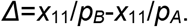

Using *x*_11_=*p*_*A*_*p*_*B*_+*D*, this becomes

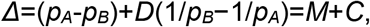

where

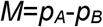

is the marginal-frequency asymmetry and

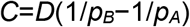

is the LD-coupled asymmetry.

This decomposition separates two sources of directional signal. The term *M* reflects asymmetry due solely to unequal allele frequencies, whereas *C* captures the additional asymmetry induced by haplotypic coupling. When *p*_*A*_≤*p*_*B*_, *M*≤0, and the sign of *C* is determined by the sign of *D*. In the rare-variant regime, the factor 1/*p*_*A*_ can amplify *C*, so that ∣*C*∣ remains large even when *D* is numerically small. This contrasts with *D*, whose feasible magnitude is bounded by allele frequencies and is therefore structurally compressed for rare variants.

The pair (*M,C*) therefore provides a directional representation of conditional carrier-set structure. In the next sections, we examine how this representation sits inside the constrained haplotype simplex and how its geometry differs from the scalar normalizations used by conventional LD measures.

## 3. Fisher geometry of the (M, D) leaf: setup

Throughout this section we fix the mean allele frequency *s*=(*p*_*A*_+*p*_*B*_)/2. Fixing *s* leaves two degrees of freedom: the frequency contrast *M*=*p*_*A*_-*p*_*B*_ and the LD coordinate *D*. We therefore study the corresponding two-dimensional (*M,D*) leaf of the haplotype simplex. The Fisher information metric on this leaf is inherited from the standard multinomial metric on the full haplotype simplex and plays a central role in the subsequent analysis.

### 3.1 Coordinates and haplotype parametrization

With p_A = s + M/2 and p_B = s − M/2, the haplotype frequencies can be written as polynomials in (s, M, D):

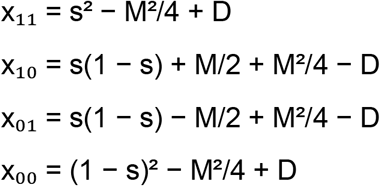

These expressions recover the expected special cases at *M*=0 and *D*=0, and define the (*M,D*) parametrization used throughout the geometric analysis below. The feasible region is obtained by imposing *x*_*ij*_≥0 for all four haplotypes.

### 3.2 Feasible domain of the (M, D) leaf

The feasible domain in (*M,D*) coordinates is determined by the non-negativity constraints *x*_11_,*x*_10_,*x*_01_,*x*_00_≥0. These constraints give four inequalities, whose equality cases correspond to the disappearance of a specific haplotype:

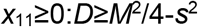

with boundary *x*_11_=0 where the *AB* haplotype vanishes;

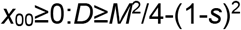

with boundary *x*_00_=0 where the *ab* haplotype vanishes;

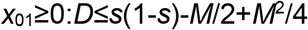

with boundary *x*_01_=0 where the *aB* haplotype vanishes;

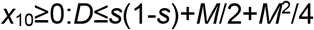

with boundary *x*_10_=0 where the *Ab* haplotype vanishes.

The marginal constraints *p*_*A*_,*p*_*B*_∈[0,1]imply ∣*M*∣≤2 *min* (*s*,1-*s*), which determines the horizontal extent of the leaf. The domain is symmetric under *M*↦-*M*, reflecting exchange of the two loci. Which lower and upper bounds are active depends on *s* and on the sign of *M*; this frequency-dependent boundary structure is analyzed in Section 4.5.

**Figure 1.**
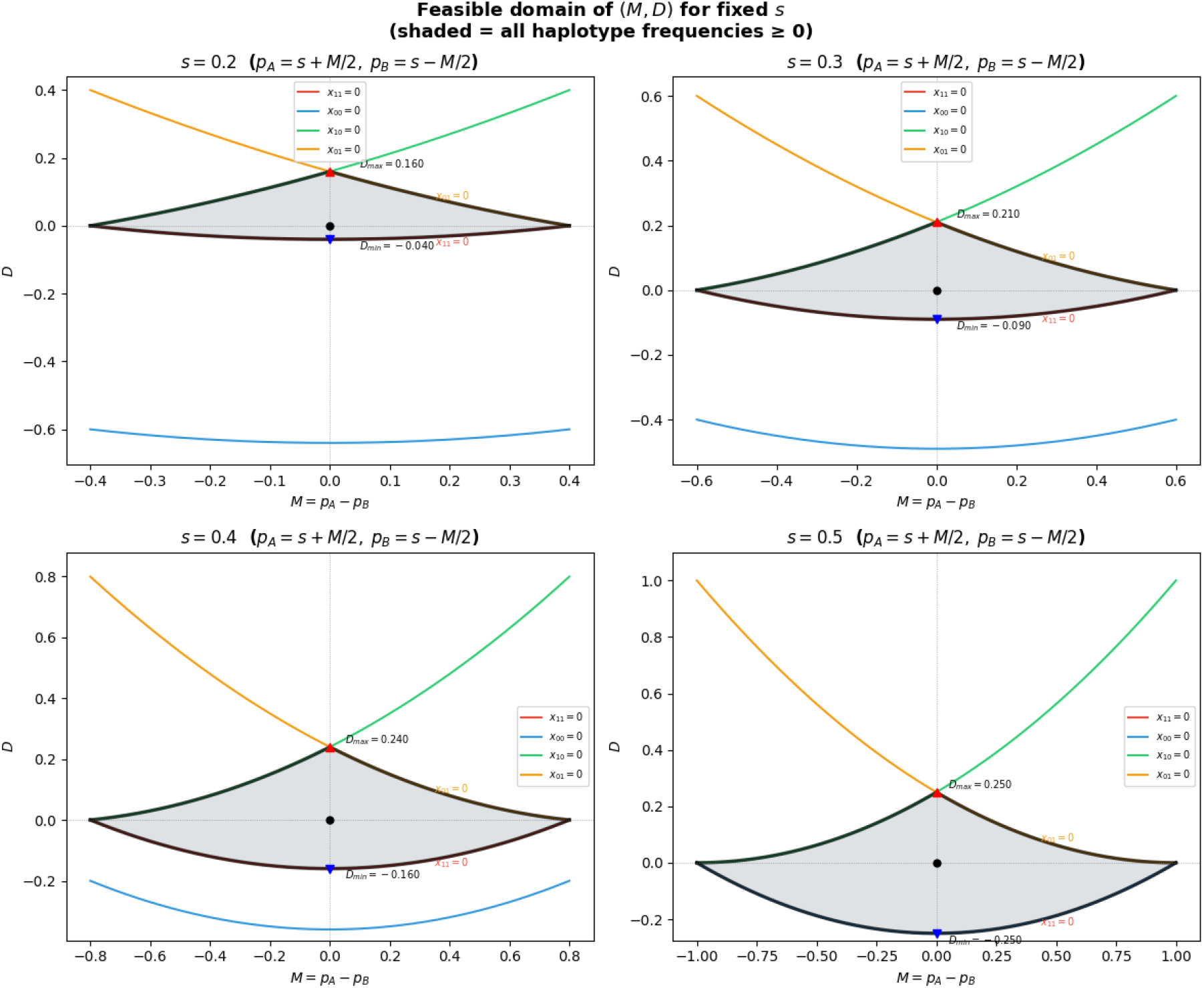
The shaded region shows the set of (*M,D*) values for which all four haplotype frequencies *x*_11_,*x*_10_,*x*_01_,*x*_00_ are non-negative, with *p*_*A*_=*s*+*M*/2 and *p*_*B*_=*s*-*M*/2. Boundary curves correspond to the disappearance of one haplotype. As *s* changes, the horizontal extent and the admissible range of *D* change, illustrating that the (*M,D*) leaf is frequency-dependent rather than a fixed Euclidean rectangle.

### 3.3 Induced Fisher metric

The induced metric follows from the multinomial Fisher form g_ij = Σ_h (∂_i x_h)(∂_j x_h) / x_h applied to the parametrization of Section 3.1; explicit expressions for the twelve partial derivatives are listed in Appendix B.

Fixing s and parameterizing the leaf by (M, D), the induced metric components are:

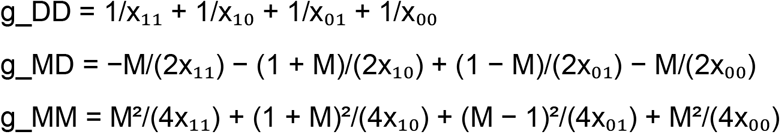

The line element on the (M, D) leaf at fixed s is dℓ^2^ = g_MM dM^2^ + 2 g_MD dM dD + g_DD dD^2^ (throughout, dℓ^2^ denotes the squared line element of the Fisher metric, with the symbol ℓ chosen to avoid collision with the mean-frequency parameter s).

## 4. Geometric structure of the (M, D) leaf

All results reported below were computed from the formulas of Section 3 and verified by independent symbolic and numerical methods. Implementation details and cross-validation procedures are summarized in Appendix B.

**Figure 2.**
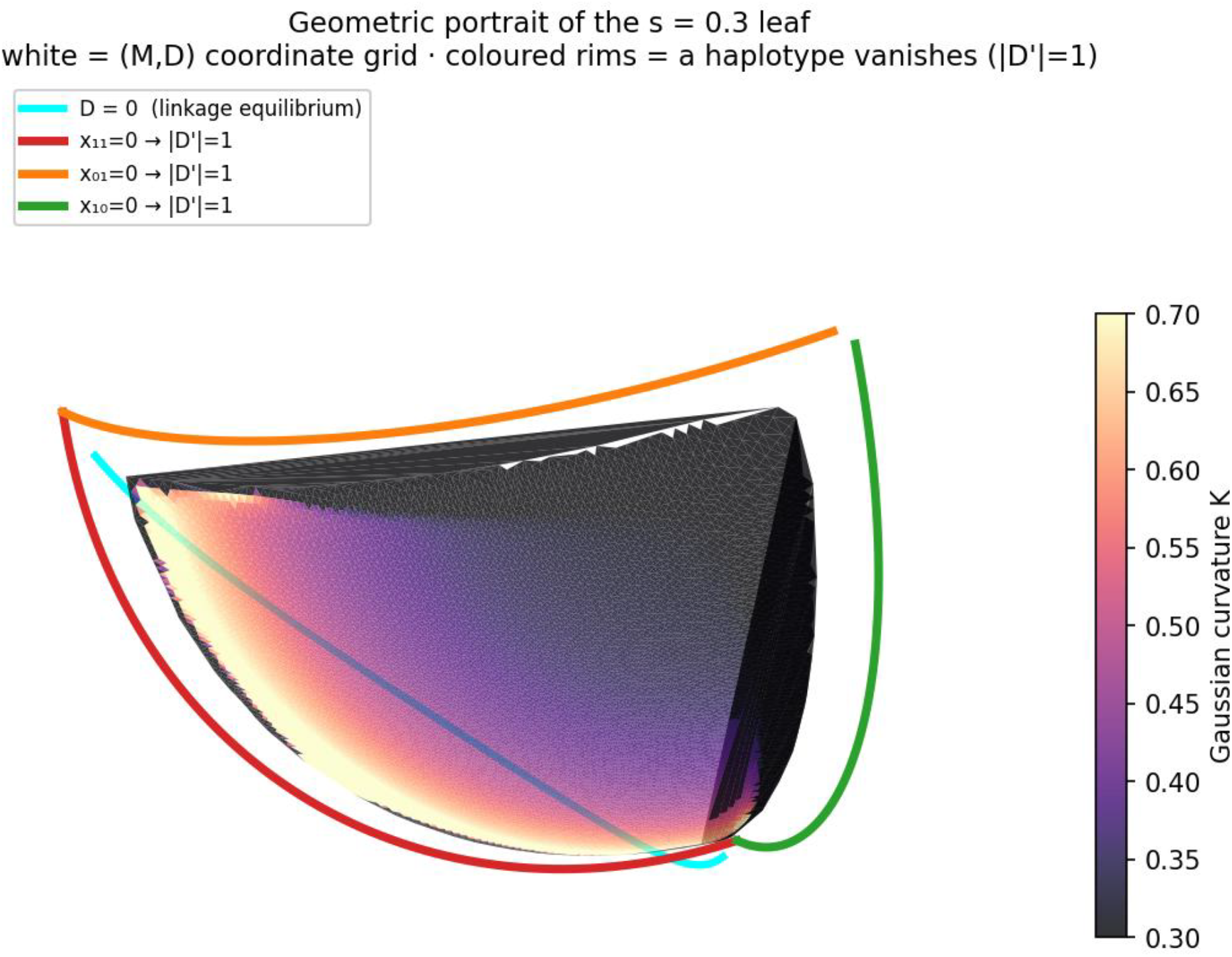
Geometric portrait of the fixed-frequency (M, D) leaf at s = 0.3. The interior is shaded by Gaussian curvature K; the white lines are the (M, D) coordinate grid. The cyan curve is the linkage-equilibrium locus D = 0, and the coloured rims mark the feasible-domain boundaries where a haplotype vanishes (|D′| = 1): x_11_ = 0 (red), x_01_ = 0 (orange), and x_10_ = 0 (green). The leaf is positively curved throughout and bounded by the four haplotype-non-negativity constraints of Section 3.2.

### 4.1 Local anisotropy at the symmetric point

At the symmetric point (s, M, D) = (1/2, 0, 0), the four haplotype frequencies are all equal:

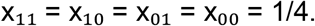

In the (M, D) coordinates, the Fisher metric at this point has components

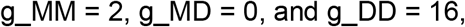

and therefore the local line element becomes

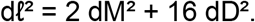

Thus, even at the maximally symmetric haplotype configuration, the metric is not isotropic in the (M, D) coordinate directions. For equal coordinate displacements, the squared Fisher length in the D direction is larger than that in the M direction by a factor of

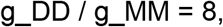

corresponding to a Fisher arc-length ratio of √8 ≈ 2.83.

This observation should not be interpreted as a new mathematical invariant, but as a local geometric property of the Fisher metric expressed in the (M, D) coordinate system. It shows that changes in LD-like displacement D and marginal imbalance M are not weighted equally by the underlying haplotype simplex. This local anisotropy provides the geometric basis for the normalization analysis in Section 6.

### 4.2 Exact orthogonality along M = 0 and D = 0

The off-diagonal metric component g_MD vanishes exactly along two loci: the symmetry axis M = 0 for all s and D, and the linkage-equilibrium axis D = 0 for all s and M. Both identities were verified by direct symbolic computation: substituting the haplotype frequencies x_h(s, M, D) into the expression for g_MD from Section 3.3 yields g_MD ≡ 0 on the set {M = 0} ∪ {D = 0}.

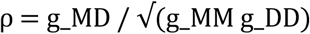

Consequently, the correlation coefficient ρ is exactly zero on each of these two loci. Non-orthogonality, ρ ≠ 0, arises only when both M ≠ 0 and D ≠ 0 simultaneously. At s = 1/2, symbolic factorization shows that ρ is proportional to the product M·D to leading order, with sign(ρ) = sign(M·D). The correlation |ρ| remains small near the origin and grows toward the corners of the feasible domain; numerical evaluation shows |ρ| reaching 0.79–0.96 near the corners, where allele-frequency asymmetry and linkage disequilibrium are simultaneously large.

The exact vanishing of g_MD along D = 0 has a natural interpretation within the Hofrichter– Jost–Tran framework. For two biallelic loci, the linkage-equilibrium manifold has a product structure: in square-root coordinates it is represented as the positive part of a product of circles, S^1^+ × S^1^+ ⊂ S^3^+, a Clifford-torus-type structure. On a fixed-s (M, D) leaf, the locus D = 0 is the corresponding linkage-equilibrium slice. Along this slice, the M direction is tangent to the linkage-equilibrium manifold, whereas the D direction points away from linkage equilibrium. The identity g_MD = 0 therefore reflects the Fisher-orthogonality between motion along the linkage-equilibrium product structure and motion away from it. Thus, the vanishing of the cross-term is not merely an isolated algebraic cancellation, but the leaf-level manifestation of the product-of-spheres geometry described by Hofrichter et al.^8^.

The two loci — M = 0, corresponding to equal marginal frequencies, and D = 0, corresponding to linkage equilibrium — therefore define natural orthogonal axes on the (M, D) leaf. Geometric coupling between marginal asymmetry and linkage disequilibrium is a higher-order effect: it is absent on either axis and becomes strongest where both deviations are simultaneously large. This structural feature plays an essential role in the interpretation of LD measures in Section 6.

**Figure 3.**
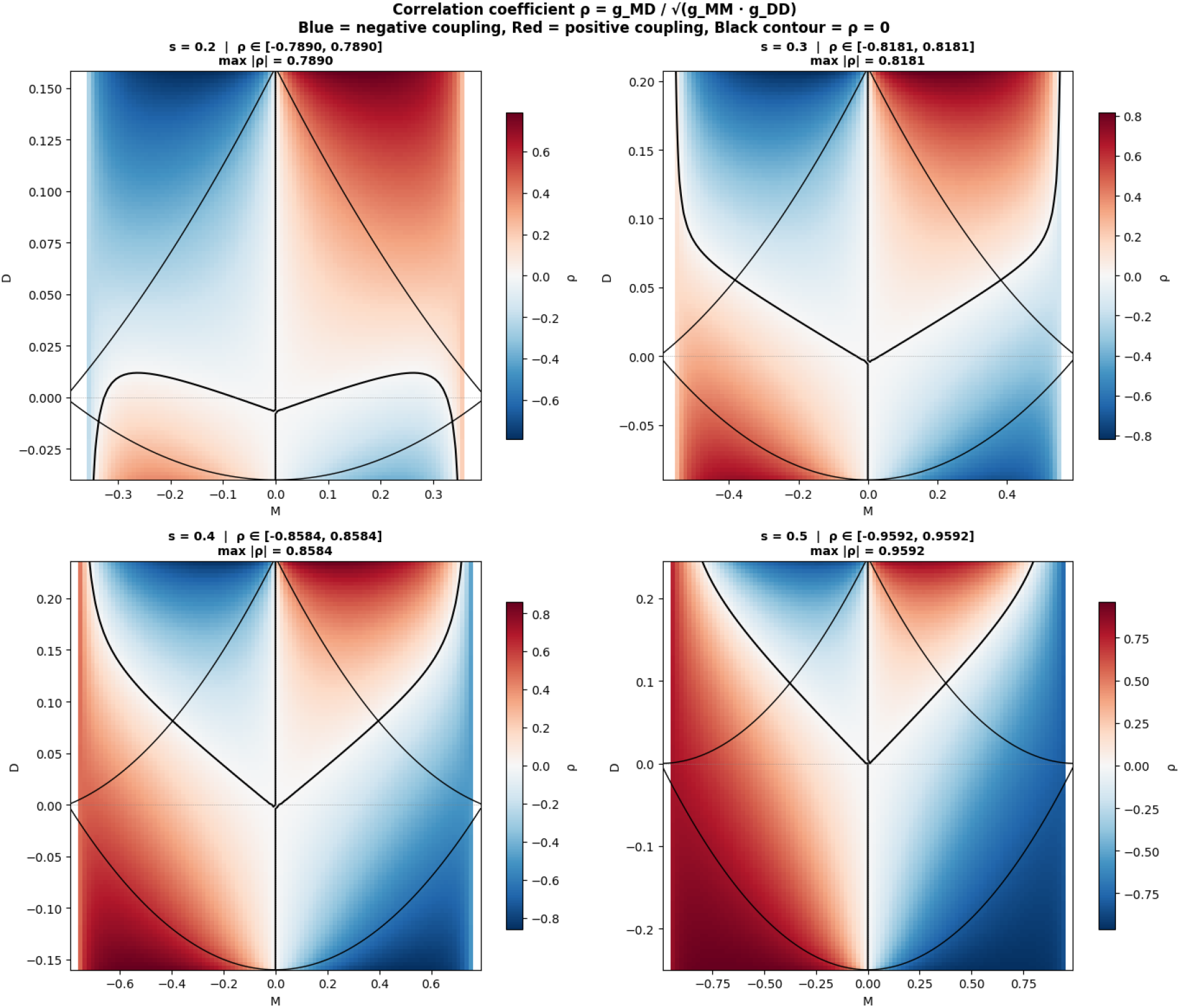
Non-orthogonality ρ = g_MD / √(g_MM g_DD) on the (M, D) leaf. ρ = 0 exactly along both M = 0 and D = 0; sign ρ = sign(MD) in the four quadrants.

### 4.3 Gaussian curvature and the rare-variant limit

The Gaussian curvature of the (M, D) leaf was computed by two independent methods: (i) the Brioschi formula applied to the metric components and their derivatives, and (ii) extrinsic curvature via the square-root embedding y_h = 2√x_h, which maps the haplotype 3-simplex isometrically onto the positive octant of the 3-sphere of radius 2 in ℝ^4^. The ambient 3-sphere has constant sectional curvature ¼, and the (M, D) leaf at fixed s is a 2-dimensional submanifold whose intrinsic Gaussian curvature is computed via the Gauss equation relating intrinsic and extrinsic curvatures. Both routes agree to numerical precision at all tested points (Appendix B).

Along the family of symmetric points {(s, 0, 0) : s ∈ (0, 1)} (to which we refer as the symmetry axis, noting that at each fixed-s leaf this is a single point), the curvature admits the exact closed form

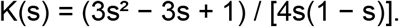

This expression can be rewritten in the Gauss-equation form

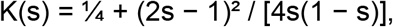

which exposes its structure directly. The leading term ¼ is the sectional curvature of the ambient 3-sphere (inherited from the Hofrichter–Jost–Tran geometry), and the second term is the non-negative correction from the extrinsic curvature of the leaf within the ambient sphere. The correction (2s − 1)^2^ / [4s(1 − s)] vanishes identically at s = ½ and is strictly positive for all other s ∈ (0, 1). This yields two immediate structural consequences.

#### First, K(s) > ¼ for all s ≠ ½

The ambient 3-sphere provides a lower bound on the leaf curvature, and the leaf is strictly more curved than the ambient manifold except at the symmetric point. Equivalently, K(s) − ¼ ≥ 0 is a sum-of-squares expression.

#### Second, at s = ½ the leaf is totally geodesic in the ambient 3-sphere

The vanishing of (2s − 1)^2^ at s = ½ corresponds, via the Gauss equation, to the vanishing of the second fundamental form of the leaf at that slice. Geometrically, the s = ½ leaf is a great 2-sphere of the ambient 3-sphere, identified by the defining relation x_11_ = x_00_. Great 2-spheres of a 3-sphere are totally geodesic submanifolds and inherit the ambient curvature, so K ≡ ¼ across the entire s = ½ leaf as a structural consequence rather than a numerical coincidence. Symbolic computation of K(M, D) at s = ½ confirms K = ¼ identically on the leaf.

For s ≠ ½, the curvature varies with position on the leaf but remains strictly positive throughout the interior in all numerical tests performed (Appendix B). The values K(s) along the symmetry axis match the closed-form formula above to at least four decimal places (Appendix B). In the rare-variant limits K(s) ∼ 1/[4s] as s → 0 and K(s) ∼ 1/[4(1 − s)] as s → 1.

**Figure 4.**
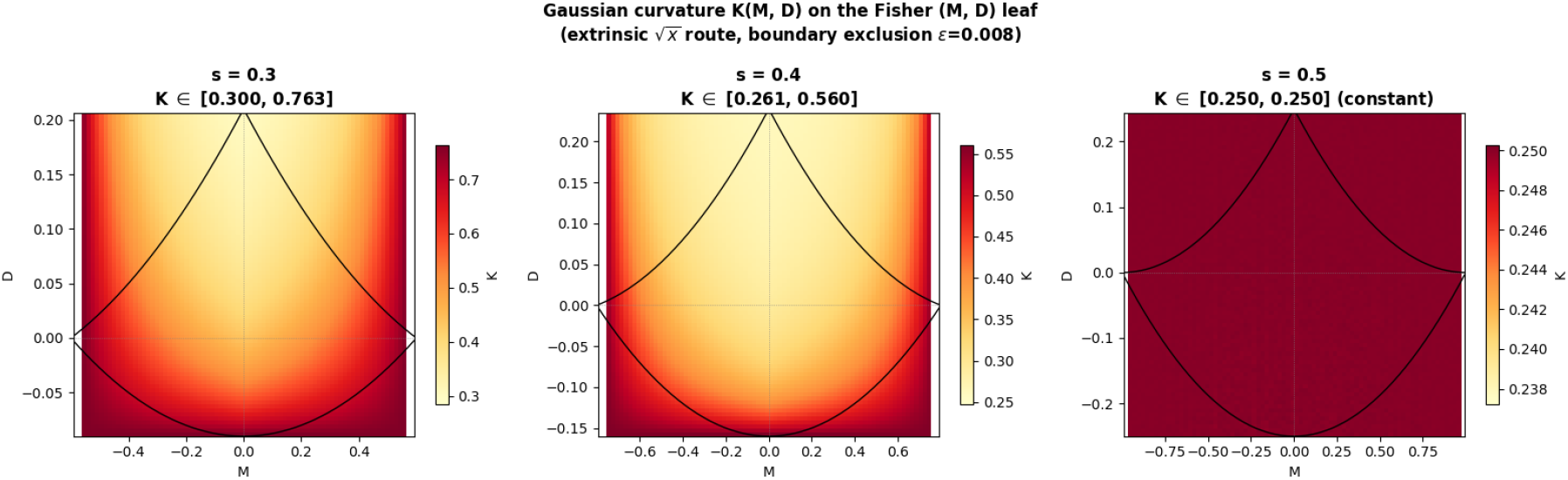
Gaussian curvature K(M, D; s) on the (M, D) leaf for s = 0.3, 0.4, 0.5, computed via the extrinsic embedding route. K is strictly positive throughout the feasible domain and becomes uniformly K = ¼ at s = ½.

**Figure 5.**
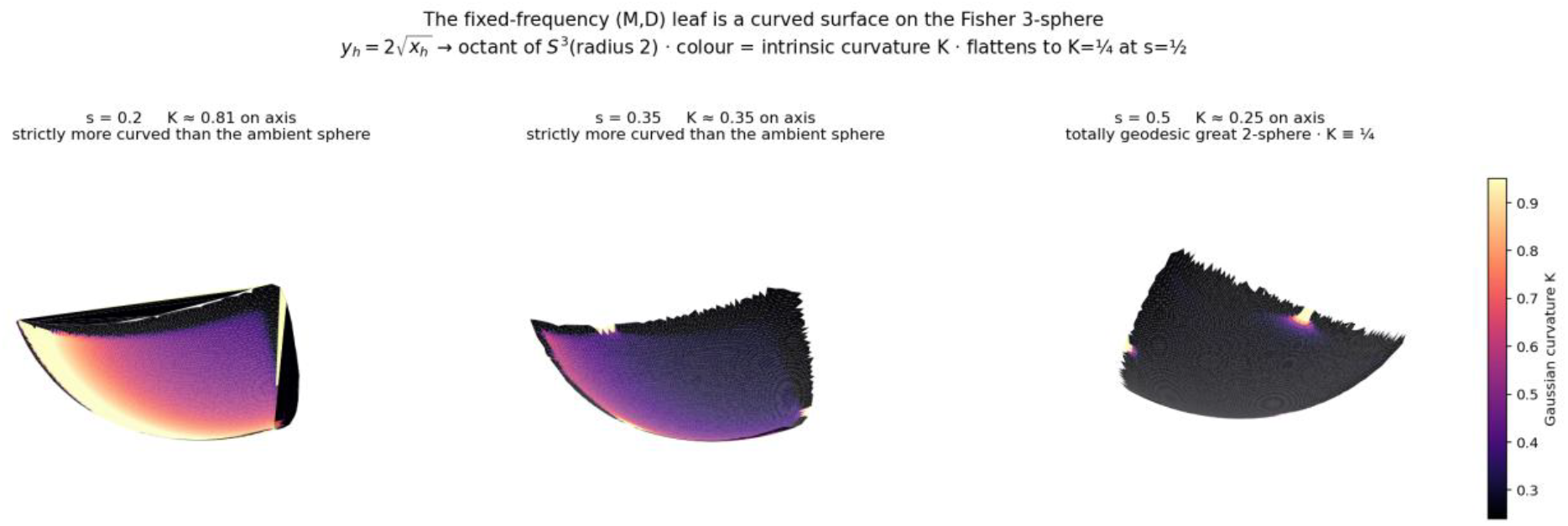
The fixed-frequency (M, D) leaf as a curved surface on the Fisher 3-sphere, shown for s = 0.2, 0.35, 0.5 under the square-root embedding y_h = 2√x_h onto the positive octant of S^3^(radius 2). Surface colour is the intrinsic Gaussian curvature K. For s ≠ ½ the leaf is strictly more curved than the ambient sphere (K > ¼); at s = ½ the leaf flattens to a totally geodesic great 2-sphere with K ≡ ¼, visualizing the Gauss-equation decomposition K(s) = ¼ + (2s − 1)^2^/[4s(1 − s)] of Section 4.3.

### 4.4 Metric volume element and boundary amplification

The metric determinant

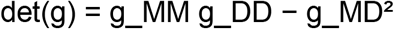

measures the local volume scaling induced by the Fisher geometry on the (M, D) leaf. For fixed s, it is moderate near the center of the feasible domain and grows rapidly near portions of the boundary where one or more haplotype frequencies approach zero. In contrast, the Gaussian curvature remains numerically stable near these fixed-s boundaries in the tested interior regions. Thus, the boundary amplification primarily reflects a divergence of metric scale, or Fisher information, rather than a curvature singularity of the leaf.

This distinction has direct statistical meaning. Near the boundary, estimation of haplotype frequencies becomes difficult because one or more haplotypes are rare or nearly absent, and the Fisher information grows accordingly. However, this increase in metric volume does not necessarily imply that the intrinsic curvature of the leaf becomes singular.

This fixed-s boundary effect should be distinguished from the separate rare-variant limit along the family of symmetry-axis points {(s, 0, 0): s in (0, 1)}. Along this family, both K(s) and det(g) diverge as s approaches 0 or 1, but with different rates. Specifically,

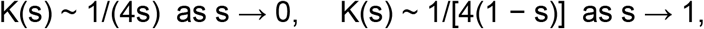

whereas

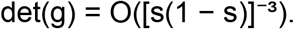

The rare-variant regime therefore combines two distinct geometric effects: strong intrinsic curvature and much stronger amplification of Fisher volume scale. Off the symmetry axis, det(g) diverges near different portions of the feasible-domain boundary through separate haplotype-frequency mechanisms, corresponding to x_h approaching zero for each of the four haplotypes in turn. We therefore do not claim a unified scaling law across the entire leaf.

**Figure 6.**
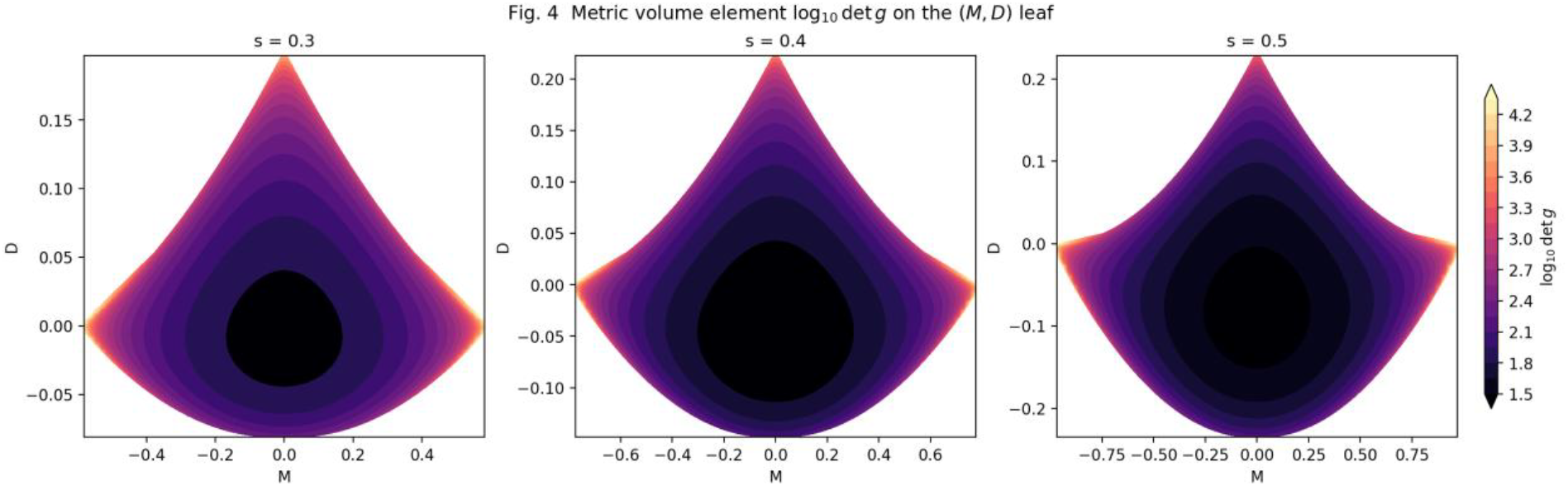
Metric volume element log_10_ det g on the (M, D) leaf, shown for s = 0.3, 0.4, 0.5. The interior is small (dark) and grows sharply toward the feasible-domain boundary where at least one haplotype frequency approaches zero. This boundary amplification reflects the divergence of Fisher information as a haplotype vanishes, and is asymmetric under D → −D because positive and negative D are constrained by different vanishing boundaries.

### 4.5 Feasible-domain asymmetry

The feasible domain introduced in Section 3.2 has a precise symmetry structure determined by the non-negativity constraints on the four haplotype frequencies. This structure is important because the apparent geometry of the (M, D) leaf is not arbitrary: it is the image of the biologically realizable haplotype simplex.

#### Symmetry under M → −M

The feasible domain is exactly symmetric under the reflection M → −M for all values of s. Under this reflection, x_11 and x_00 are invariant because they depend on M only through M^2^, whereas x_10 and x_01 are exchanged:

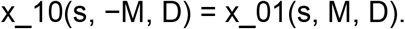

The four non-negativity constraints are therefore permuted among themselves, and the feasible domain is preserved. This reflects the exchange symmetry between the two loci.

#### Asymmetry under D → −D

In contrast, the feasible domain is not generally symmetric under the reflection D → −D. Positive and negative values of D are constrained by different haplotype-vanishing boundaries. Positive D is limited by the disappearance of the repulsion haplotypes x_10 or x_01, whereas negative D is limited by the disappearance of the coupling haplotypes x_11 or x_00. As a result, the admissible positive- and negative-D ranges are generally unequal.

At s = 1/2, the leaf attains its highest internal symmetry: the coupling haplotypes satisfy x_11 = x_00, and the geometry simplifies substantially. However, this does not imply full invariance of the feasible domain under D → −D for arbitrary M. Even at s = 1/2, the upper D boundary is determined by x_10 and x_01, whose constraints depend on |M|, whereas the lower boundary is determined by x_11 = x_00. Exact symmetry of the positive and negative D ranges occurs only at the central point s = 1/2, M = 0. Away from this point, the imbalance between the upper and lower D bounds is a direct consequence of haplotype non-negativity constraints.

#### Dependence on s

As s decreases toward 0, or symmetrically increases toward 1, the feasible domain shrinks and becomes increasingly constrained by boundary effects. In the rare-variant regime, the allowed range of D is strongly limited because small changes in haplotype configuration can drive one of the four haplotype frequencies to zero. Each boundary has a direct biological interpretation: it represents the disappearance of one haplotype. Thus, the feasible-domain asymmetry is the geometric signature of biological realizability constraints in the two-locus haplotype simplex.

## 5. The (M, C) representation

The geometric results of Sections 3 and 4 were formulated on the (M, D) leaf, where D is a smooth coordinate. For the purposes of LD interpretation, however, the (M, C) plane offers a more transparent visualization. We introduce it here as a representation of the leaf, not as a new global coordinate system.

The decomposition

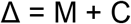

is most naturally displayed in the (M, C) plane. Here,

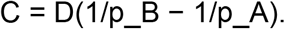

However, the change of variable from (M, D) to (M, C) is singular on the line {M = 0}. Along this line, p_A = p_B, and every value of D maps to C = 0. Therefore, (M, C) fails to parameterize the leaf globally.

Away from M = 0, the map (M, D) → (M, C) is locally smooth, and results established on the (M, D) leaf can be re-expressed in the (M, C) plane for visualization. The advantage of this representation is that Δ = constant corresponds to a straight diagonal line, making the decomposition Δ = M + C visually transparent. All quantitative claims in this paper are formulated on the (M, D) leaf; the (M, C) displays below are used for intuition only.

**Figure 7.**
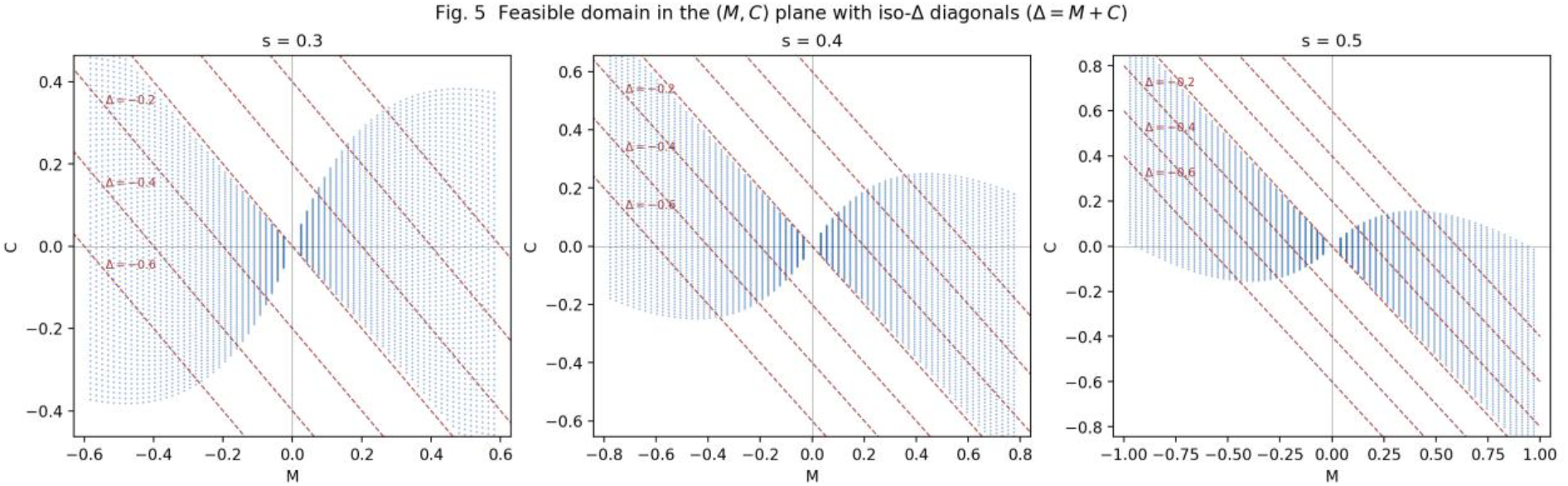
The feasible domain in the (M, C) plane for s = 0.3, 0.4, 0.5, with iso-Δ diagonals Δ = M + C shown in red. Along M = 0 the change of variable (M, D) → (M, C) is singular: all D values map to C = 0. As s → 1/2, the domain becomes more symmetric under M → −M. Each red diagonal is a level set of the conditional-probability asymmetry Δ = P(A | B) − P(B | A).

**Figure 8.**
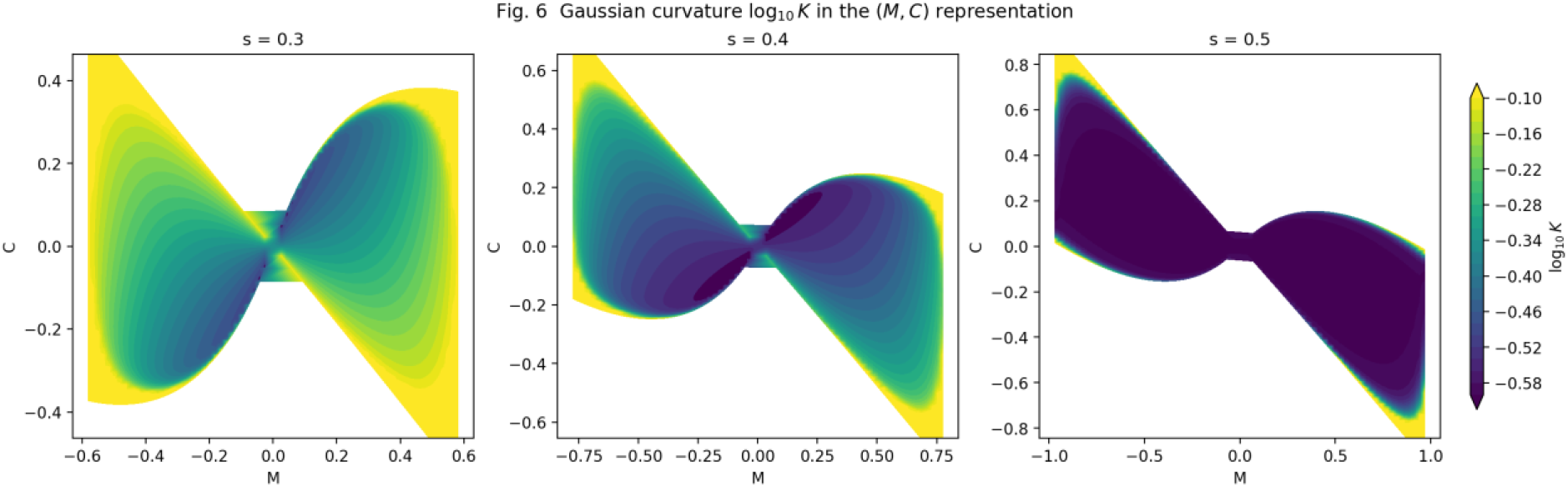
Gaussian curvature log_10_ K displayed in the (M, C) representation for s = 0.3, 0.4, 0.5. At s = 1/2, K ≡ 1/4 uniformly across the leaf, as predicted by the totally-geodesic great 2-sphere result of Section 4.3; log_10_ K is therefore a single dark value. For s ≠ 1/2, K exceeds 1/4 and grows toward the feasible-domain corners, illustrating positive curvature enhanced away from the symmetric point.

**Figure 9.**
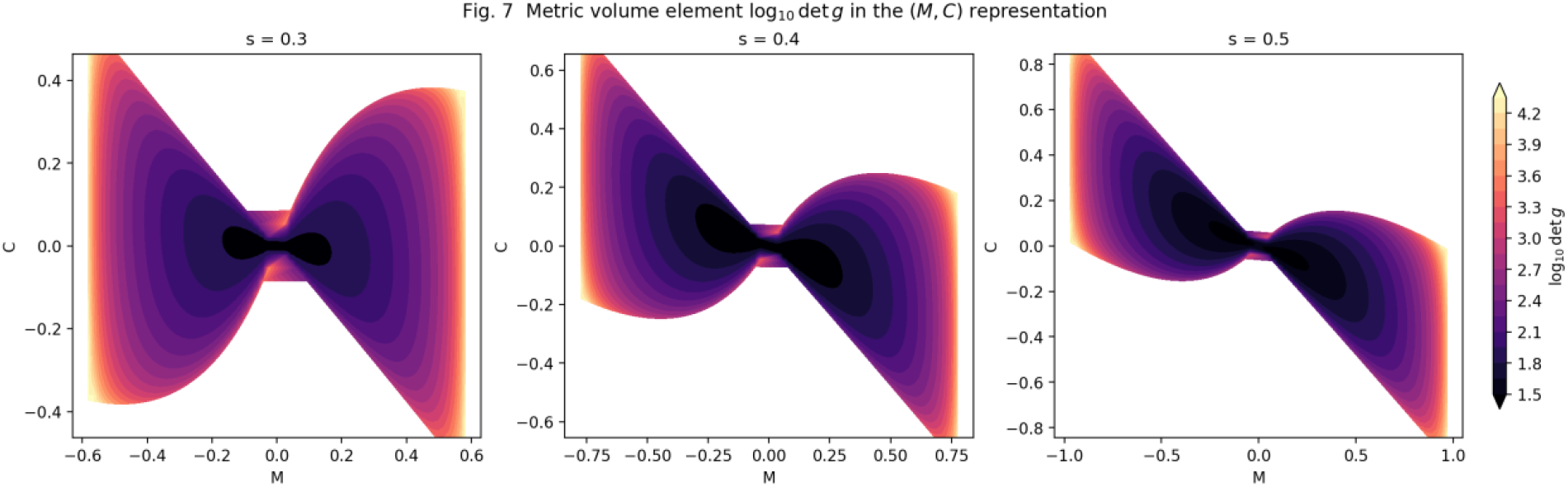
Metric volume element log_10_ det g in the (M, C) representation for s = 0.3, 0.4, 0.5. The interior is small near the linkage-equilibrium locus and diverges toward the feasible-domain boundary. The (M, C) view makes boundary amplification visually parallel to Figure 6 while exposing the iso-Δ direction along which the decomposition Δ = M + C is naturally read.

**Figure 10.**
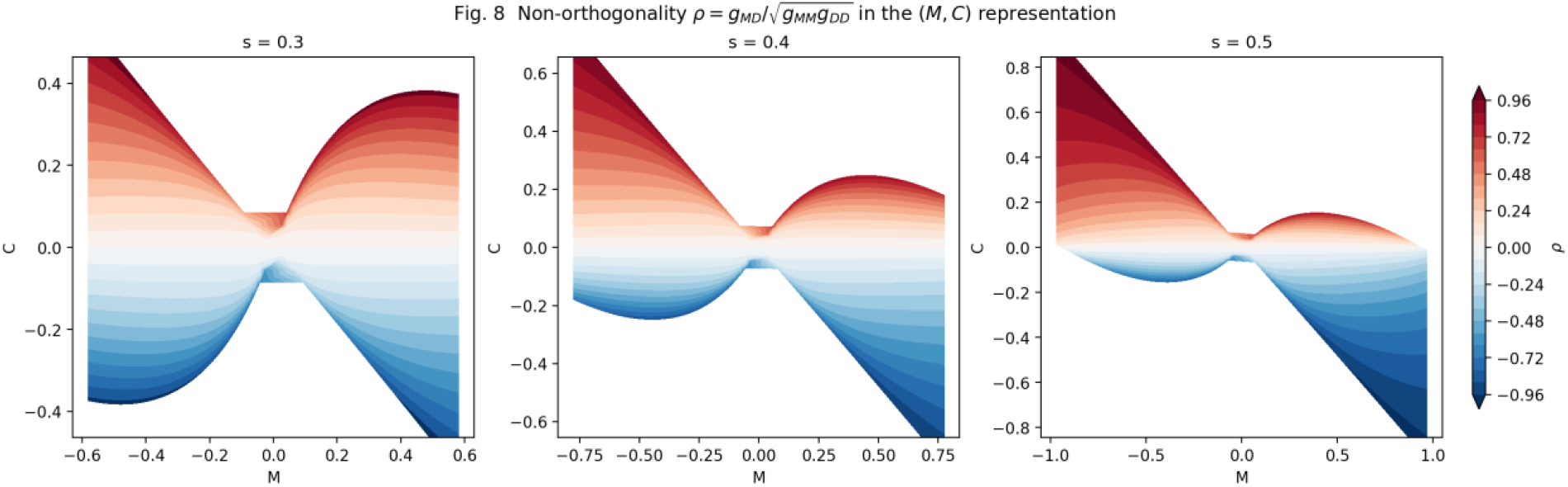
Non-orthogonality ρ = g_MD / √(g_MM g_DD) in the (M, C) representation for s = 0.3, 0.4, 0.5. The horizontal white band at C = 0 corresponds to D = 0, along which ρ vanishes exactly by the orthogonality result of Section 4.2. ρ approaches +1 in corners where M and D have the same sign, and −1 where they differ, matching the |ρ| ≈ 0.79–0.96 near-corner values reported in the text.

## 6. Standard LD measures as scalar functions on the Fisher-curved simplex

### 6.1 The interpretational problem and the Euclidean-treatment mismatch

Multiple statistics have been proposed to quantify linkage disequilibrium, each defined as a different frequency-dependent scalar function on the haplotype 3-simplex (Lewontin 1964; Hedrick 1987; Kang and Rosenberg 2020). These scalars — D, D’, r^2^, |C| — are widely understood to capture “different aspects” of LD, and are routinely used in downstream analysis pipelines as if they were Euclidean quantities: thresholded at fixed values, compared across loci and populations, binned into ranges, and ranked by magnitude.

Consider a concrete example. Let two biallelic loci be in complete LD (D’ = 1) with allele frequencies p_A = 0.01 and p_B = 0.30, giving D = p_A(1 − p_B) = 0.007. The standard LD measures report D = 0.007, D’ = 1.000, r^2^ = 0.024, and |C| = 0.68. These are not approximation errors; each value is correct given its definition. Yet they yield mutually inconsistent biological conclusions about the same pair of loci in the same population.

The question is not which measure is right but why correct measures disagree, and — more operationally — what structures become invisible when these scalars are used in the Euclidean manner that current practice assumes. The core observation of this paper is that the haplotype 3-simplex, as characterized in Sections 3 and 4, is not Euclidean: it is a positively curved Riemannian manifold with anisotropic metric, frequency-dependent curvature divergence, and frequency-asymmetric feasible domain. Scalar functions on this curved manifold, when treated as Euclidean quantities, lose information in regions where the geometric structure most strongly deviates from the Euclidean ideal. Sections 6.2 and 6.3 identify the specific measures and regimes where this information loss is most severe.

### 6.2 Standard LD measures as frequency-dependent normalizations of D

#### Two-axis classification of LD measures

The scalar measures considered below can be organized along two independent design axes. The first is the *symmetry axis*: symmetric measures treat the two loci as exchangeable, whereas asymmetric measures retain the directional labelling of the two loci. The second is the *evaluation axis*: sign-preserving or directional measures retain the coupling phase or the direction of conditional-probability asymmetry, whereas magnitude-only measures collapse the signal to a non-negative quantity.

Under this classification, signed D and signed D′ are symmetric and sign-preserving. r^2^, |D′|, and D^2^-based normalizations are symmetric and magnitude-only: r^2^ is a squared correlation, whereas |D′| preserves boundary saturation but discards sign. |C| is asymmetric in construction but magnitude-only, while signed C and Δ = P(A|B) − P(B|A) — whose decomposition Δ = M + C is introduced in Section 2 — are asymmetric and sign-preserving / directional. ALD-type measures occupy the asymmetric magnitude family, although for biallelic SNP pairs ALD reduces to r^2^ and therefore adds no SNP-pair-level resolution beyond r^2^.

Geometrically, magnitude evaluation compresses directional information much as the modulus |z|, or squared modulus |z|^2^, compresses a complex number z: the sign of D is eliminated before summation, and the two-component structure of the conditional probability pair (P(A|B), P(B|A)) collapses to a single non-negative magnitude. Sign-preserving asymmetric measures retain this two-component structure and are the natural scalar functions to place on the Fisher-curved simplex when directional information is to be preserved. The remainder of this section treats each measure in turn and identifies the regimes in which Euclidean treatment produces systematic distortion.

The standard LD measures are distinct frequency-dependent normalizations of the coefficient D, each defined as a scalar function of the haplotype frequencies:

#### D = x_11_ − p_A p_B

The LD coefficient is bounded by |D| ≤ min(p_A q_B, p_B q_A), so when p_A is small, |D| is structurally suppressed regardless of haplotype association strength. D compresses the rare-variant signal.

#### D′ = D / D_max

Here D_max denotes the sign-specific feasible bound. Normalization by D_max removes the upper-bound constraint: the signed normalization D′ saturates at ±1 at the feasible boundary, whereas |D′| saturates at 1. In the rare–common regime, even modest association drives |D′| to its maximum. |D′| inflates the rare-variant signal as a boundary-saturation measure, while signed D′ additionally preserves coupling phase.

#### r^2^ = D^2^ / (p_A q_A p_B q_B)

The squared correlation divides by the product of heterozygosities and, crucially, squares D, eliminating the sign and collapsing coupling-phase (D > 0) and repulsion-phase (D < 0) relationships into a single non-negative value. When p_A ≪ p_B, the factor p_A q_A ≈ p_A constrains r^2^ ≤ p_A / p_B ≪ 1 regardless of haplotype association. r^2^ compresses the rare–common signal and discards directional information.

#### |C| = |D| · |1/p_B − 1/p_A|

The coupling component C scales |D| by the reciprocal-frequency difference. When p_A ≪ p_B, the factor 1/p_A dominates, amplifying |C| far beyond |D|. |C| amplifies the rare-variant signal; the signed component C preserves the direction of the conditional-probability asymmetry, whereas |C| itself is used only as a magnitude.

From an information-geometric perspective, the normalization in r^2^ = D^2^ / (p_A q_A p_B q_B) can be read as a variance normalization under linkage equilibrium — the denominator is the asymptotic variance of D under the independence hypothesis — and equivalently as a Fisher-information weighting of the squared deviation from independence. This helps explain two of r^2^’s characteristic behaviors. First, r^2^ is symmetric under the exchange of the two loci because the weighting treats both marginals identically. Second, r^2^ systematically suppresses rare–common associations: the same weighting that makes r^2^ distance-like around linkage equilibrium also compresses configurations in which one allele is rare. By contrast, the coupling term C = D · (1/p_B − 1/p_A) applies a directional, one-sided frequency weighting to D, preserving asymmetry that r^2^ removes through squaring. In this sense, r^2^ and C represent two contrasting ways of reading the same underlying LD coefficient: r^2^ as a symmetric distance-like deviation from equilibrium, C as a directional asymmetry in the conditional probabilities P(A|B) and P(B|A) that — as introduced in Section 2 — is aligned with carrier-set inclusion between the two variants.

The pattern runs from compression to amplification:

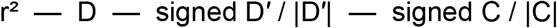

r^2^: ceiling ≤ p_A/p_B; sign discarded

D: bounded by p_A; sign preserved

signed D′: sign preserved and boundary-normalized

|D′|: boundary saturation preserved, sign discarded

signed C: directional asymmetry preserved

|C|: rare-variant magnitude amplified

Each measure faithfully reports the value of its own normalization. The disagreements arise because different normalizations of D, evaluated as scalar functions on the same Fisher-curved manifold, respond differently to the underlying geometric structure.

#### Dual Fisher-information reading of r^2^ and C

The symmetric Fisher-information reading of r^2^ admits a complementary directional reading for C. The Bernoulli Fisher information at allele frequency p is I(p) = 1/[p(1 − p)] = 1/p + 1/(1 − p), the sum of a positive-outcome branch 1/p (the information carried by observing the allele present) and a negative-outcome branch 1/(1 − p) (the information carried by observing it absent). The r^2^ denominator p_A q_A p_B q_B is the product of the reciprocals of these two full Bernoulli Fisher informations, so that r^2^ = D^2^ · I(p_A) · I(p_B) weights D^2^ symmetrically by the two full Bernoulli Fisher informations. The coupling term C = D·(1/p_B − 1/p_A) uses only the positive-outcome branch 1/p at each locus and takes their difference, weighting D antisymmetrically by the gradient of positive-branch Fisher information between the two loci.

The two measures are thus dual faces of the same Fisher structure rather than alternatives chosen by convention. r^2^ is a symmetric, distance-like reading: it asks how far the joint haplotype distribution sits from independence, weighted isotropically by the information each locus carries about the presence or absence of its allele. C is an antisymmetric, directional reading: it asks in which direction the conditional probabilities P(A∣B) and P(B∣A) diverge, weighted by the imbalance in positive-branch information between the two loci. This duality explains the rare–common regime behavior directly. Where one allele is rare, the symmetric weighting in r^2^ is dominated by the vanishing heterozygosity p(1 − p) → 0, which, combined with the compression of D by the same factor, yields the structural ceiling r^2^ ≤ p_A q_B / (q_A p_B) derived above. The antisymmetric weighting in C is dominated instead by the one-sided divergence 1/p → ∞, which amplifies the signal. Suppression and amplification are not two distinct pathologies but two readings of the same curved geometry along different axes of symmetry.

### 6.3 Geometric sources of disagreement: where Euclidean treatment distorts

The exact results of Section 4 identify three geometric features of the Fisher metric on the (M, D) leaf under which Euclidean treatment of scalar measures produces systematic distortion. For each feature, the Fisher geometry predicts both the regime in which distortion is strongest and the direction in which each measure fails.

i. **Metric anisotropy**. At the symmetric point (s, M, D) = (½, 0, 0), the line element is dℓ^2^ = 2 dM^2^ + 16 dD^2^, so that g_DD / g_MM = 8 in metric-tensor units. A unit Euclidean displacement in D carries eight times the squared Fisher information of a unit Euclidean displacement in M. Measures that apply a uniform Euclidean threshold to D (or to a scalar function of D) therefore impose different effective Fisher-distance thresholds depending on the direction in which the threshold is crossed. The anisotropy varies across the leaf, and is most pronounced when s ≠ ½ or when the leaf is probed near feasible-domain boundaries.
ii. **Positive Gaussian curvature, intensifying toward rare-variant limits**. Along the symmetry axis, the curvature K(s) = ¼ + (2s − 1)^2^ / [4s(1 − s)] is strictly positive, attains its minimum ¼ at s = ½, and diverges as s → 0 or s → 1. The rare-variant regime is not only metrically stretched but intrinsically more curved. Under Euclidean treatment, a scalar measure’s behavior calibrated in the near-flat balanced regime will fail to capture its behavior as curvature diverges: the mapping between Euclidean value and Fisher-geometric content is not preserved across the leaf, and the divergence is concentrated precisely where rare-variant pairs are located.
iii. **Asymmetry of the feasible domain**. The domain is exactly symmetric under M ↦ −M but asymmetric in D whenever s ≠ ½. Measures that treat D symmetrically (e.g., r^2^ = D^2^/pq) discard this intrinsic asymmetry of the space on which they are evaluated. Euclidean binning of such measures treats regions of different feasible-domain extent as comparable, producing imbalanced statistical power across D > 0 and D < 0 configurations.

### 6.4 Non-orthogonality and coordinate coupling

The exact orthogonality result of Section 4.2 — g_MD = 0 precisely on M = 0 and D = 0, with ρ proportional to M · D elsewhere — has a direct interpretational consequence. The geometric coupling between marginal asymmetry and linkage disequilibrium is a higher-order effect: it vanishes identically when either M = 0 (equal allele frequencies) or D = 0 (linkage equilibrium) and is most pronounced when both deviations are simultaneously large. This is precisely the rare–common, high-LD regime where biological signal often concentrates and where LD measures disagree most strongly.

Measures that ignore the g_MD term — for example, by treating M and D as independent Euclidean coordinates — introduce distortion proportional to the local non-orthogonality. Along the two exact orthogonality loci, such decoupled treatments are exact; off these loci, the distortion grows with |MD|. The Fisher-geometric analysis thus identifies a natural coordinate system in which the anisotropy of the metric cleanly separates into pure-M and pure-D contributions on the orthogonality loci {M = 0} and {D = 0}, with the leading off-axis cross-term scaling as |MD| near the symmetric slice s = ½. The full g_MD at arbitrary (s, M, D) is not captured by this leading form.

### 6.5 Boundary singularity, mixed populations, and the Wahlund regime

The divergence of det(g) near the feasible-domain boundary has a direct statistical interpretation: points near the boundary, where one or more haplotype frequencies approach zero, are information-geometrically far from the interior even when they appear close in Euclidean (M, C) coordinates. This is the geometric manifestation of the statistical difficulty of estimating haplotype frequencies near zero.

As established in Section 4.3, the metric divergence det(g) ∼ 1/[s(1 − s)]^3^ and the curvature divergence K(s) ∼ 1/[s(1 − s)] both concentrate in the rare-variant regime, though with different rates. The rare-variant region thus combines two distinct geometric effects — large Fisher information (cubically diverging) and strong intrinsic curvature (linearly diverging) — both pushing measures away from their balanced-regime behavior under Euclidean treatment.

A complementary regime in which the Fisher geometry predicts systematic Euclidean distortion is the mixed-population (Wahlund) regime. When a sample is pooled across subpopulations with different allele frequencies, D picks up a between-subpopulation component. Writing w_k for normalized subpopulation weights (Σ_k w_k = 1), D_k for the within-subpopulation LD, and p_{A,k}, p_{B,k} for subpopulation-specific allele frequencies, the pooled coefficient is D_pool = Σ_k w_k D_k + D_between, where the second term is the weighted covariance of the subpopulation allele frequencies, D_between = Σ_k w_k p_{A,k} p_{B,k} − (Σ_k w_k p_{A,k})(Σ_k w_k p_{B,k}); equivalently, in pairwise form, D_between = ½ Σ_{k,k’} w_k w_{k’} (p_{A,k} − p_{A,k’})(p_{B,k} − p_{B,k’}). This component inflates D relative to its within-subpopulation value, and hence inflates r^2^, even when within-subpopulation LD vanishes.

Because r^2^ inherits this inflation through squared D, r^2^ in pooled cohorts cannot separate genuine within-population LD from spurious inflation driven by subpopulation structure. The decomposition Δ = M + C makes the pooled-sample frequency imbalance visible alongside C: a nonzero M flags that the marginals differ, so a frequency-driven contribution is less easily mistaken for a linkage signal. This is disclosure, not correction — the Wahlund inflation itself rides on D (and hence on C), while M only reflects the pooled marginal contrast, so actual separation of within-from between-subpopulation LD requires population-stratified computation — whereas r^2^ folds the pooled imbalance into the squared numerator where it cannot be inspected at all.

More generally, Fisher-metric-weighted analogues of common LD-based summary statistics would be more robust to allele-frequency composition than their Euclidean counterparts, down-weighting boundary regions and high-curvature regimes without ad hoc thresholding. The qualitative signature (systematic MAF-dependent divergence of r^2^ and |C|, systematic inflation of r^2^ in mixed cohorts) is confirmed in the companion paper^2^.

### 6.6 Relation to prior work

The connection between the Fisher metric and population genetics has been recognized since Shahshahani (1979). Hofrichter, Jost, and Tran (2019) developed the full geometric framework for recombination dynamics, establishing that the haplotype 3-simplex under the Fisher metric is the positive octant of the 3-sphere of radius 2 (sectional curvature ¼), that linkage equilibria correspond to products of spheres (specifically, a Clifford-torus structure for two biallelic loci), and that the Ohta–Kimura formula for LD decay admits a clean geometric interpretation on this manifold.

The present work uses this established geometry as the setting in which standard LD measures r^2^, D, D’, and |C| are re-examined. The ambient sectional curvature ¼ appears as the leading term in our Gauss-equation decomposition K(s) = ¼ + (2s − 1)^2^/[4s(1 − s)], and the Clifford-torus linkage-equilibrium manifold of Hofrichter et al. manifests at the leaf level as the exact vanishing of g_MD along D = 0 (Section 4.2). Our contribution is not to re-derive the ambient manifold but to characterize the (M, D) leaf at fixed mean frequency s — its induced metric, curvature, and feasible-domain structure — and to identify how standard scalar measures project onto this leaf.

In a complementary direction, Kang and Rosenberg (2020)^9^ systematically characterized the mathematical properties of five LD statistics defined as normalizations of D, and Tanaka ^6^ proposed an equal-tempered LD measure derived from a geometric interpretation of r^2^. These contributions document the frequency-dependent behavior of individual measures but do not provide a unified account of why the measures disagree. The present work connects the Shahshahani / Hofrichter et al. line (Fisher geometry of population genetics) and the Kang– Rosenberg / Tanaka line (catalogue and reformulation of LD measures) by showing that the measure discrepancies catalogued in the latter arise from the geometric structure established in the former.

### 6.7 Implications

This geometric perspective has three operational consequences.

**First**, the question of which LD measure to use is not answerable by a single choice: r^2^ is well-behaved in the common–common corner where metric and domain are nearly isotropic, while |C| retains signal in the rare–common regime where both depart from isotropy. The geometry dictates which measure is faithful in which region, and the correspondence is deterministic, not empirical.

**Second**, the ceiling r^2^ ≤ p_A q_B / (q_A p_B) is a structural bound on rare–common pairs, not a finite-sample artifact: r^2^-based GWAS tagging cannot detect haplotype associations that generate directional LD signals in this regime, and no increase in sample size can lift the bound.

**Third**, directional measures such as Δ, signed C, and max(P(B|A), P(A|B)) preserve the directional (carrier-set) asymmetry that quadratic normalizations discard. The companion paper (Ichikawa, preprint) confirms empirically that these measures recover nested haplotype structure within r^2^ bins where r^2^ itself performs at chance.

## 7. Discussion

On the Hofrichter–Jost–Tran Fisher-curved simplex, standard LD measures r^2^, D, D’, and |C| are scalar functions whose disagreements are controlled by the leaf-level geometric structure characterized in Sections 3–4. Five observations are worth highlighting.

### Relation to the Hofrichter–Jost–Tran geometry

The Fisher-curved simplex was established by Hofrichter et al. (2019) from Wright–Fisher recombination dynamics. The Gauss-equation decomposition K(s) = ¼ + (2s − 1)^2^/[4s(1 − s)] makes the relationship explicit: the ambient 3-sphere curvature ¼ is the leading term, and the leaf correction is a sum of squares vanishing exactly at s = ½, where the leaf is totally geodesic as a great 2-sphere of the ambient 3-sphere. The linkage-equilibrium Clifford torus of Hofrichter et al. appears at the leaf level as the exact vanishing of g_MD along D = 0. The present work takes this geometry as established and characterizes how the standard scalar measures behave on the (M, D) leaf.

### Euclidean treatment of scalar LD measures produces systematic distortion

r^2^, D, D’, and |C| are scalar functions on the Fisher-curved simplex. Current practice thresholds, bins, and compares these scalars as Euclidean quantities. The Fisher metric predicts that this mismatch produces distortion concentrated in three regimes: rare–common allele-frequency combinations (where curvature diverges and the feasible domain is most asymmetric), boundary-proximal configurations (where det(g) diverges), and mixed-population samples (where Wahlund-type inflation enters D and is masked by squaring in r^2^). The geometric prediction is sharp: within a fixed r^2^ bin, r^2^ should carry no information about nested carrier-set structure, whereas the boundary-preserving magnitude |D′| should retain it — |D′| = 1 marks exactly the vanishing of a haplotype, i.e. the nested / complete-LD boundary — because that information is present in the haplotype frequencies but compressed out by the r^2^ normalization. This prediction is tested and confirmed in phased 1000 Genomes data in the companion paper.

### The rare–common blind spot is structural, not statistical

The bound r^2^ ≤ p_A q_B / (q_A p_B) is a theorem of the r^2^ normalization, not a finite-sample artifact. No increase in sample size can lift the ceiling. Association signals that are accessible under |C|, max(P(B|A), P(A|B)), or other directional measures can be invisible under r^2^ regardless of statistical power. For rare-variant GWAS tagging, fine-mapping, and imputation reference selection, this has direct operational consequences: reliance on r^2^ alone imposes a frequency-dependent blind spot that should be addressed by including directional measures in routine reporting.

### Sign preservation carries information that squaring discards

r^2^ discards the sign of D by squaring, collapsing coupling and repulsion phases. Within a fixed r^2^ bin, two kinds of information that r^2^ discards remain recoverable: the boundary-preserving magnitude |D′| retains the nested / complete-LD saturation (|D′| = 1 when a haplotype is absent), and the signed directional measures — signed C and Δ — retain the coupling-versus-repulsion phase. Both are predicted to discriminate nested structure while r^2^ is at chance, a prediction borne out empirically in the companion paper^2^. The sign and direction of the conditional-probability asymmetry — information that squaring eliminates — therefore carry genuine haplotype-structure content. Downstream tasks that depend on distinguishing subclade structure, coupling versus repulsion phase, or between-subpopulation versus within-subpopulation LD are candidates where sign-preserving measures should be inspected alongside r^2^.

### Mixed-population LD analysis requires explicit frequency decomposition

In pooled cohorts, D picks up a Wahlund between-subpopulation component that inflates r^2^ through squaring. The decomposition Δ = M + C does not, by itself, separate within-population from between-population LD: M is a property of the pooled marginals and does not perform a stratified variance decomposition, and strict separation of the Wahlund component requires population-stratified computation. What the decomposition does provide is a display of the pooled-sample frequency imbalance alongside Δ: M contributes directly to the conditional-probability asymmetry whenever p_A ≠ p_B, so inspecting (M, C) rather than r^2^ alone makes it harder to mistake a frequency-driven contribution for a linkage signal. This is disclosure, not correction. For cross-population LD comparison and cross-ancestry meta-analysis, population-stratified computation remains the necessary step; the decomposition simply makes the pooled-sample frequency component visible in the same representation that displays the directional asymmetry.

### A unifying observation

The three empirical phenomena most often encountered in LD practice — the r^2^ ceiling in the rare–common regime, the amplification of ∣C∣ at rare variants, and the Wahlund inflation of r^2^ in mixed populations — have traditionally been treated as separate methodological caveats to be handled case by case. On the Fisher-curved simplex they are three visible consequences of a single geometric fact: variant pairs occupy different regions of a curved manifold whose metric, curvature, and feasible-domain shape all vary with allele frequency, so that the same algebraic change in D has different geometric meaning depending on where on the manifold it is evaluated. Euclidean treatment of scalar LD measures implicitly assumes a flat geometry in which algebraic and geometric differences coincide; the distortions encountered in practice are precisely the signature of the mismatch between this assumption and the intrinsic Fisher geometry characterized in Sections 3–4.

A final remark concerns the scope of the framework. The geometric structure characterized here is intrinsic to the haplotype 3-simplex and therefore applies to any pair of biallelic loci, regardless of the organism, population history, or trait context in which LD is studied. The empirical confirmation in the companion paper, using human 1000 Genomes data^1^, is illustrative. Extensions to multi-allelic loci and higher-dimensional haplotype spaces are natural and follow the general product-of-spheres framework of Hofrichter et al.^8^, but are beyond the present scope.

## 8. Implementation note

All symbolic results were verified in SymPy with independent cross-checks. Numerical results on the feasible domain and along the symmetry axis used grid evaluations at N ∈ {60, 70, 80} per axis, finite-difference step sizes h ∈ {10^−7^, 10^−6^, 10^−5^, 10^−4^}, and boundary exclusion thresholds ε ∈ {0.005, 0.02, 0.05, 0.10}. Reproducibility materials are available from the author on request.

## Acknowledgments

No acknowledgments.

## Author Contributions

Y.I. conceived the study, developed the mathematical and geometric analysis, performed the symbolic and numerical verification, interpreted the results, and wrote the manuscript.

## Use of Artificial Intelligence Tools

Large language model tools were used to assist with manuscript drafting, language editing, code review, and organization of the presentation. All mathematical derivations, computational analyses, interpretations, and final statements were checked and approved by the author. The author takes full responsibility for the content of the manuscript.

## Data Availability

No new empirical datasets were generated in this study. The empirical analyses using 1000 Genomes Phase 3 data that test the geometric predictions of this note are reported in the companion paper.

## Code Availability

Symbolic verification scripts and numerical curvature-evaluation code used to reproduce the results of Sections 3–4 are available in the fisher_geometry/ subdirectory of the unified project repository at https://github.com/mountbook-lab/directional-ld-mc, archived at Zenodo as release v0.1.1 with DOI https://doi.org/10.5281/zenodo.21098988. These include scripts for evaluating the induced Fisher metric on the (M, D) leaf, computing Gaussian curvature by the Brioschi-formula route, computing curvature through the square-root embedding route, and generating the geometric figures of this note. The same repository hosts the code for the empirical and Wright–Fisher analyses of the companion paper (Ichikawa 2026, bioRxiv 2026.07.01.735767) under the dynamic_topology/ and empirical_1000g/ subdirectories, together with supplementary animations under animations/.

## Appendix B. Partial derivatives and verification of the Gaussian curvature

### B.0 Partial derivatives of the haplotype frequencies

With haplotype frequencies parameterized as in Section 3.1, the twelve partial derivatives with respect to (s, M, D) are: ∂x_11_/∂s = 2s, ∂x_10_/∂s = ∂x_01_/∂s = 1 − 2s, ∂x_00_/∂s = 2(s − 1); ∂x_11_/∂M = ∂x_00_/∂M = −M/2, ∂x_10_/∂M = (1 + M)/2, ∂x_01_/∂M = (M − 1)/2; and ∂x_11_/∂D = ∂x_00_/∂D = +1, ∂x_10_/∂D = ∂x_01_/∂D = −1. All twelve were verified by independent symbolic computation and are used in the derivation of g_MM, g_MD, and g_DD given in Section 3.3.

The Gaussian curvature of the (M, D) leaf was verified by two independent numerical routes together with symbolic computation.

#### B.1 Route I: Brioschi formula

The first route applies the Brioschi formula to g_MM, g_MD, g_DD and their derivatives, computed by finite differences on a grid covering the feasible domain. Grid resolutions N = 60, 70, 80 per axis, step sizes h ∈ {10^−7^, 10^−6^, 10^−5^, 10^−4^}, and boundary exclusion thresholds ε ∈ {0.005, 0.02, 0.05, 0.10} were tested.

#### B.2 Route II: Extrinsic curvature via the square-root embedding

The second route uses y_h = 2√x_h, mapping the haplotype 3-simplex onto the positive octant of the 3-sphere of radius 2 in ℝ^4^ under the Fisher metric. The ambient 3-sphere has constant sectional curvature ¼. The Gaussian curvature of the 2-dimensional (M, D) leaf embedded in this 3-sphere is computed from the Gauss equation relating intrinsic and extrinsic curvatures, giving K_intrinsic = ¼ + det(II)/det(g), where II is the second fundamental form of the leaf in the ambient sphere. The Gauss-equation form of K(s) in Section 4.2 is the direct consequence of this decomposition.

#### B.3 Route III: Symbolic computation along the symmetry axis and at s = ½

Along (M, D) = (0, 0) for general s, SymPy-based symbolic computation of the Brioschi formula yields the exact closed form K(s) = (3s^2^ − 3s + 1)/[4s(1 − s)] = ¼ + (2s − 1)^2^/[4s(1 − s)]. At s = ½, symbolic computation on (M, D) yields K = ¼ identically. This constant-curvature property reflects the fact that the s = ½ leaf is the great 2-sphere x_11_ = x_00_ of the ambient 3-sphere, hence totally geodesic and inheriting the ambient curvature.

#### B.4 Cross-validation

At all tested (s, M, D) configurations, Routes I and II agree to within 0.01 in the interior (ε ≥ 0.02). Along the symmetry axis, both routes match the closed-form formula from Route III to at least four decimal places across s ∈ {0.1, 0.2, …, 0.9}. Examples: K(0.1) = 2.0278, K(0.2) = 0.8125, K(0.3) = 0.4405, K(0.4) = 0.2917, K(0.5) = 0.2500. The symmetry K(s) = K(1 − s) was verified numerically.

**Figure B1.**
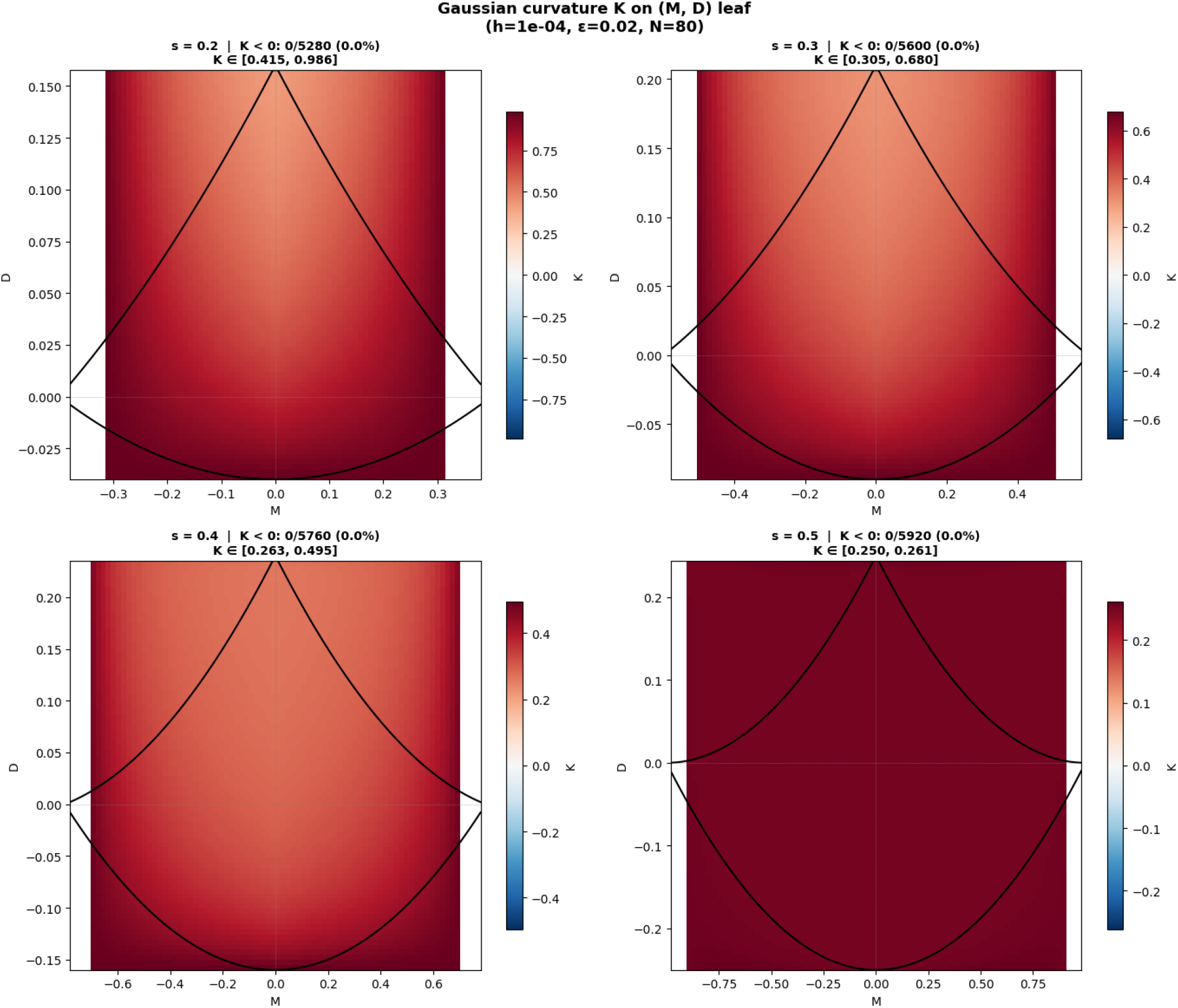
Gaussian curvature K on the (M, D) leaf computed via Route I (Brioschi formula on g_MM, g_MD, g_DD with finite-difference derivatives) at s ∈ {0.2, 0.3, 0.4, 0.5}. Parameters: h = 10^−4^, ε = 0.02, N = 80 per axis. K > 0 at every grid point (0/N_tot_ negative across all panels), and K → ¼ uniformly at s = ½.

**Figure B2.**
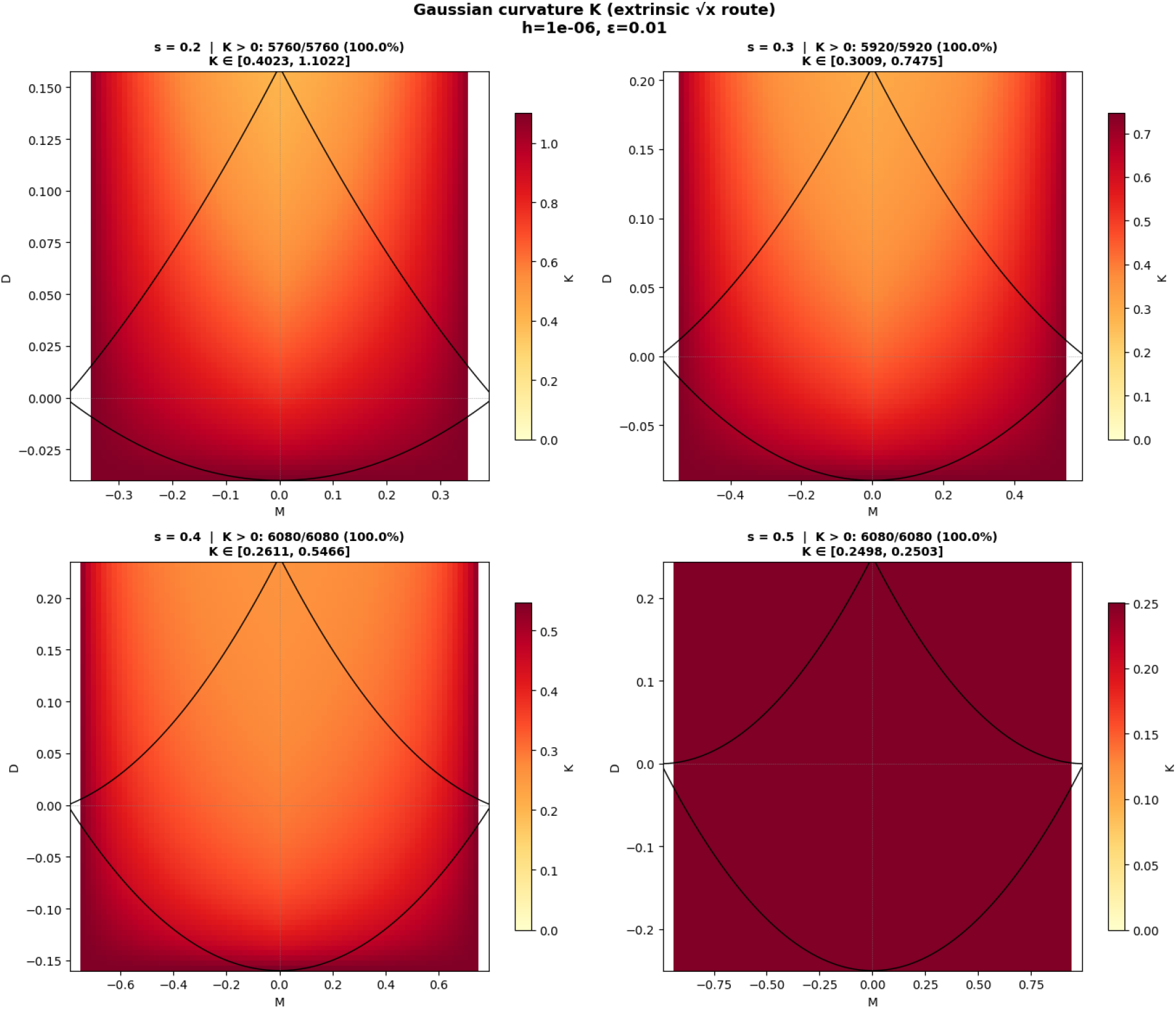
Gaussian curvature K on the (M, D) leaf computed via Route II (extrinsic curvature through the y_h = 2√x_h embedding) at the same s values. Parameters: h = 10^−6^, ε = 0.01. K > 0 at every grid point (100% across all four panels), and K → ¼ at s = ½. Numerical agreement between Routes I and II (Figures B1 and B2) is consistent across the leaf and supports the closed-form result of Section 4.3.

#### B.5 An excluded route

A fourth approach — direct symbolic computation via Christoffel symbols and the Riemann tensor — was initially attempted but produced spurious zero curvature due to incomplete symbolic simplification of complex rational expressions in SymPy. Once this computational artifact was identified (by comparison with Routes I–III, which consistently yielded K > 0), the Christoffel-based route was excluded from the final analysis. This experience underscores the value of cross-validation across structurally different computational methods.

#### B.6 Robustness

Across 64 combinations of (N, h, ε) × {s = 0.2, 0.3, 0.4, 0.5} tested via Routes I and II, K > 0 was observed at every grid point in the interior, with no exception. The sign of K was invariant under all parameter variations.

